# A study of the neuronal injury biomarkers pNF-H and UCHL1 in serum, CSF and urine in a cohort of thoracic endovascular aortic repair (TEVAR) patients

**DOI:** 10.1101/2021.03.02.433405

**Authors:** Adam Beck, Vedrana Marin, Jay Deng, Irina Madorsky, Bin Ren, Yichen Guo, Dan Neal, Gerry Shaw

## Abstract

A collection of longitudinal serum, cerebrospinal fluid (CSF) and urine samples were collected from a cohort of 50 patients undergoing thoracic endovascular aortic repairs (TEVAR). Samples were taken multiple time per day out to 5 days post operation and were probed with novel electrochemiluminescent assays specific for the phosphorylated axonal form of the major neurofilament subunit NF-H (pNF-H) and for ubiquitin C-terminal hydrolase 1 (UCHL1). Control blood samples showed small signals for pNF-H and in some cases rather larger signals for UCHL1. The presence of UCHL1 in these control blood samples was convincingly verified by western blotting with multiple well characterized UCHL1 antibodies and by mass spectroscopy. Elevated levels of both pNF-H and UCHL1 in blood and CSF in recovering TEVAR patients were associated with poorer outcomes. In particular release of UCHL1 into blood over several hours following TEVAR and peaking at any time over 1 ng/ml was a very strong predictor of patient death and was associated with renal failure and spinal cord ischemia (SCI). An unexpected finding was that high levels of UCHL1 were detected in certain urine samples, again in association with SCI, renal failure and poor patient outcome. We also present epitope mapping data on the UCHL1 monoclonal antibodies used including data on the widely used commercially available MCA-BH7. These studies suggest that measurement of the levels of both proteins in the blood, CSF and urine of TEVAR patients may be of clinical utility. However this study also raises questions about the origin and significance of UCHL1 both in control blood, in patient blood samples and in urine.

## Introduction

Surgeons perform thousands of thoracic endovascular aortic repair (TEVAR) operations each year in the United States to treat aortic aneurysms, traumatic aortic transections, and aortic dissections. TEVAR often involves coverage of important collateral blood flow to the spinal cord, including the left subclavian and intercostal arteries. Coverage of these collaterals leads to clinically detectable spinal cord ischemia (SCI) in 3-10% of patients [1–3]. Preoperative clinical characteristics can be predictive of SCI, and include the extent of aorta to be covered, male gender, presence of an infrarenal abdominal aortic aneurysm (AAA), previous AAA repair, and pre-existing renal insufficiency [4–8]. Unfortunately, these characteristics are not sensitive or specific for the development of SCI, leading to little clinical guidance in the peri-operative setting.

Early interventions to treat SCI after TEVAR are available, and include lumbar cerebrospinal fluid (CSF) drainage for cord decompression [9], pharmacologic elevation of blood pressure [10], treatment with free radical scavengers [11], and with steroids [10]. Currently, the physical exam is the only mechanism for diagnosing SCI in patients after TEVAR, which is an unreliable and often grossly ineffective measure. The development of SCI has a significant impact on patients’ post-operative outcomes. Amabile et al. found that paraplegia after TEVAR was associated with a nearly ten-fold increase in the risk for in hospital mortality [6]. Additionally, data from European studies have shown that the overall survival of patients with SCI was 25% at 12-months, while patients without SCI had an 80% 12-month survival [12]. Although SCI may be a surrogate for other untoward complications leading to death in these patients, prediction and prevention of SCI may improve long-term outcomes. It follows that if SCI can be identified in its early stages and an intervention undertaken, great potential benefits may exist for the patient. Recent studies have shown that various proteins derived from the cells of the CNS may be detected in CSF and blood following a variety of CNS compromises. Investigators at our institution as well as others have discovered biomarkers of CNS injury in both brain [13–18] and spinal cord injury [19, 20]. The levels of these markers may reflect the timing, location and severity of injury, allowing the identification of windows for therapeutic intervention prior to the development of symptoms, the assessment of prognosis, and the monitoring of responses to therapy.

Two well studied biomarker proteins are the phosphorylated axonal form of the major neurofilament subunit NF-H (pNF-H) [21–24] and cytoplasmic enzyme ubiquitin C-terminal hydrolase 1 (UCHL1) [25, 26]. The pNF-H protein is one of the components of neurofilaments, structural proteins found only in neurons. Since the pNF-H form of NF-H is normally only found in axons, detection of significant amounts of this protein in CSF, blood or other fluids points to recent axonal damage or degeneration [27, 28]. This is of great potential utility as axons are unusually sensitive to mechanical and metabolic stress [29, 30]. The pNF-H protein has an unusual multi-epitope nature that allow it to be detected in CSF and blood using antibody body based assays unusually efficiently [13]. UCHL1 is an abundant cytoplasmic enzyme heavily expressed by neurons, although, as discussed here, it is expressed ubiquitously at lower levels in other tissues. It may also leak into blood and CSF following CNS injury and degeneration and elevated levels of this protein in both CSF and blood have been associated with poor patient outcome [17, 31]. UCHL1 was originally named Protein Gene Product 9.5 (Pgp9.5), identified as a major protein spot on stained 2 dimensional gels of brain tissue extracts which was not obvious on similar gels of non CNS extracts [32]. Antibody to Pgp9.5 was and still is widely used to identify neurons, their processes and certain cell types. It was many years before the enzymatic function of this protein was discovered, leading to the renaming of Pgp9.5 as UCHL1 [33]. UCHL1 cleaves ubiquitin monomers from other proteins and is important to allow ubiquitin to be recycled but also to release ubiquitin from polyubiqutin and ubiquitin fusion proteins encoded in the genome. The majority of UCHL1 in the CNS is clearly in gray matter, while the majority of pNF-H is found in white matter, so we have proposed that the ratio of these two proteins may provide information about not only the severity but also the location of a CNS lesion [31].

There have been relatively few studies of CNS injury biomarkers following SCI [34] and virtually none following TEVAR. Here we briefly describe novel and highly sensitive assays for these two proteins and show how these assays were used to monitor CSF, blood and in some cases urine of a cohort of TEVAR patients.

## Materials and Methods

### Patients

The sampling protocol was approved by the institutional research board (IRB) of the University of Florida and 50 TEVAR patients were enrolled at the Shands Hospital, Gainesville, Florida. Patients ranged in age from 36.2 to 85.2 years, with the average being 71.5 and the median being 74.6. 34 were male (69.4%). A single patient treated emergently for a contained rupture of a thoracoabdominal aneurysm died during the 5 day sampling period and 7 more had expired within one year of the operation, while the remainder recovered to varying degrees (Table 1). A total of 529 serum samples were taken and all were analyzed for UCHL1 and pNF-H. The IRB allowed one sample to be taken preoperatively, one at the time of the operation, one every 4 hours out to 24 hours and then one every 6 hours out to 120 hours. There was some variance from the protocol for various unforeseen circumstances although in most cases the full protocol was accomplished. In the case of 9 patients blood samples were also collected at a 30 day follow up. Blood was collected in 10mL Vacutainer tubes (BD catalog # 366430) with no anticoagulants and silicone coated interiors. All samples were collected in the operating room, inpatient space or outpatient clinic space. After collection, blood sample tubes were inverted several times, kept at room temperature in an upright position for several minutes to allow clotting, then were placed in an ice bath for 10-15 minutes. They were spun at 3,000 rpm for 15 minutes in a clinical centrifuge and separated into ~1 mL aliquots. If samples were collected after hours and could not be processed immediately, they were placed on ice at the patient’s bedside until research staff were available. After centrifugation and separation into aliquots, the samples were immediately placed in storage boxes in a −80°C freezer. A total of 293 CSF samples were obtained by lumbar puncture from most patients, although due to low volume CSF output along with for staffing and other organizational reasons not every blood sample had a corresponding CSF sample. Following a modification in the original IRB, urine samples were taken in the case of 15 patients allowing the collection of 108 urine samples. A urine sample was generally taken preoperatively, at zero hours and, when possible, at the same times as the blood and CSF samples. Both CSF and urine samples were collected in the same type of Vacutainer tube as the blood samples and were processed in parallel and in the same fashion as the blood samples. 20 control serum samples were obtained from BioreclamationIVT (Nassau, New York). 10 were from males, the average age was 41.1 and the range was 21 to 61.

**Table 1:** Patient and procedure characteristics and outcomes (N=50)

|  |  |
| --- | --- |
| **Complications** |  |
| Stroke | 2 (4.0) |
| Myocardial infarction, | 2 (4.0) |
| Gastrointestinal hemorrhage | 1 (2.0) |
| Acute renal failure | 1 (2.0) |
| Respiratory failure | 1 (2.0) |
| Spinal cord ischemia | 8 (16.0) |
| Reintubation | 3 (6.0) |
| Renal failure | 2 (4.0) |
| Pulmonary failure | 1 (2.0) |
| Ischemia | 3 (6.0) |
| Paraplegia | 1 (2.0) |
| Death | 3 (6.0) |
| Cerebrovascular accident | 1 (2.0) |
| **Procedure characteristics** |  |
| **Spinal drain** |  |
| None | 4 (8.0) |
| Post-implant | 1 (2.0) |
| Pre-implant | 45 (90.0) |
| Fluoro | 44.2 (49.1); 21.5 [10, 62] (0, 200) |
| Contrast | 79.8 (60.9); 72.5 [30, 121] (0, 250) |
| Procedure time (minutes) | 222 (166); 186 [97, 281] (45, 900) |
| Estimated blood loss | 432 (873); 200 [100, 300] (0, 4200) |
| **Discharge characteristics** |  |
| Length of stay | 10.8 (12.1); 6 [4.25, 13] (2, 60) |
| **Disposition** |  |
| Home | 25 (50.0) |
| Home w/family help | 6 (12.0) |
| Home w/home care | 4 (8.0) |
| Inpatient facility | 12 (24.0) |
| Died in house | 3 (6.0) |
| **Ambulation at discharge** |  |
| Ambulating | 28 (58.3) |
| Ambulating with assistance | 11 (22.9) |
| Non-ambulatory | 3 (6.2) |
| Bedridden | 2 (4.2) |
| Unknown | 1 (2.1) |
| NA (died) | 3 (6.2) |
| **Ambulation at 1 month** |  |
| Ambulating | 31 (63.3) |
| Ambulating with assistance | 2 (4.1) |
| Non-ambulatory | 1 (2.0) |
| Bedridden | 0 (0) |
| Unknown | 10 (20.4) |
| NA (died) | 5 (10.2) |
| **Ambulation at 6 months** |  |
| Ambulating | 19 (38.8) |
| Ambulating with assistance | 1 (2.0) |
| Non-ambulatory | 1 (2.0) |
| Bedridden | 0 (0) |
| Unknown | 21 (42.9) |
| NA (died) | 7 (14.3) |
| **Ambulation at 12 months** |  |
| Ambulating | 9 (18.8) |
| Ambulating with assistance | 1 (2.1) |
| Non-ambulatory | 0 (0) |
| Bedridden | 0 (0) |
| Unknown | 29 (60.4) |
| NA (died) | 9 (18.8) |

Patient outcome was assessed on several numerical scales. Disposition at discharge was assessed as 0 at home, 1 = at home with family help, 2 = at home with home care, 3 = in an inpatient facility and 4 = died in house. 25 patients were in category 0, 6 in category 1, 4 in category 2, 11 in category 3 and 3 in category 4. We graded ambulation at discharge and at 1, 6 and 12 month follow up in the following categories; 0 = ambulating, 1 = ambulating with assistance, 2 = non ambulatory, 3 = bedridden, 4 = unknown, 5 = dead.

### Electrochemiluminescent pNF-H and UCHL1 assays

We developed assays optimized for the Mesoscale Discovery (MSD) electrochemiluminescence (ECL) platform based on ELISAs described previously [18, 27, 35]. For the pNF-H MSD assay we used affinity purified mouse monoclonal anti-pNF-H for antigen capture and antigen affinity purified chicken polyclonal anti-pNF-H for detection. The detection antibody was directly conjugated with MSD sulfotag NHS ester (MSD catalog R91AN-1) following the standard manufacturer protocol at a molar ratio of tag reagent to antibody of 20:1. All experiments were performed on MSD standard 96 well plates (MSD catalog L15XA). In brief, plates were coated overnight at 4°C with 30 μL of purified capture antibody diluted in phosphate buffered saline (PBS) at pH=7.5 at a final concentration of 2 μg/mL. Plates were washed twice with PBS and incubated in 100 μL of 3% BSA in PBS for 1 hour at room temperature (RT) on a microplate shaker at 500 rpm. Plates were then washed with 100 μL TBSt (Tris buffered saline plus 0.1% Tween-20) on the microplate washer. Samples were run in duplicate with 25 μL of sample in each well. The serum samples were run at dilutions of 1:1 while CSF and urine were run at 1:4, in both cases in sample diluent. Samples were incubated for 1 hr at RT with vigorous shaking as before. Protein standards were applied as serial dilutions of purified bovine pNF-H standard starting at 5 ng/mL in 25 μL sample diluent. After incubation plates were washed four times with TBSt on a plate washer, followed by incubating with 25 μL detection antibody prepared in 0.1% BSA in TBSt at a final concentration of 1 μg/mL for 1 hr at RT with vigorous shaking. Finally, the plates were washed five times with TBSt on a plate washer, 150 μL of MSD read buffer T (MSD catalog R92TC-1) was added to each well and the plates were immediately placed on a MSD Sector Imager 2400 for imaging and data collection. MSD Discovery Workbench 3.0.18 software was used to calculate protein standard concentrations and normalized results with in-plate controls to obtain the final concentration of each sample. The antibodies and protein standards are commercially available from EnCor Biotechnology Inc.

The UCHL1 assay is based on the regular ELISA described in Lewis et al. and uses the same antibody pair [36]. MSD plates were coated with affinity purified mouse monoclonal anti-UCHL1 at 2 μg/mL in PBS and detection was performed with affinity purified and sulfotagged rabbit polyclonal anti-UCHL1. Labeling was performed using a 20:1 molar ratio of tag to antibody with the MSD sulfotag reagent using the MSD protocol. Plates were coated with 30 μL per well of purified capture antibody diluted in PBS at a final concentration of 2 μg/mL overnight at 4°C. Washing and incubation was performed as for the pNF-H MSD assay, though samples were made 1:3 in diluent, and the detection antibody was used at a concentration of 0.5 μg/mL. The protein standard was recombinant human His-tagged UCHL1 expressed in and purified from *E. coli*. The antibodies and protein standards are commercially available from EnCor Biotechnology Inc. The detailed characteristics of both the pNF-H and UCHL1 MSD assays will be described in a forthcoming publication (Shaw et al. in preparation). Western blotting on the urine samples shown in Figure 4 was performed by standard procedures using the same two UCHL1 antibodies as the MSD assay.

### Biochemical and proteomic studies

Rat and mouse tissues were homogenized and sonicated in 50 mM Tris-HCl, 5 mM EDTA, 100 mM NaCl, 1% SDS, pH = 8.0 plus Pierce protease inhibitor (Life Technologies Cat 88666) and were centrifuged at 13,000g to remove insoluble material. Samples were then boiled and run out on SDS-PAGE gels for western blotting. Control human blood samples were processed to remove the most abundant proteins using albumin and IgG depletion SpinTrap columns (GE Healthcare Cat 28-9480-20) following the manufacturers instructions. Western blotting was performed using standard methods and fluorescent blot imaging was performed on the LiCor Odyssey machine. Certain control blood samples depleted of IgG and albumin were run out on SDS-PAGE gels, stained with Coomassie brilliant blue and the 24 kDa region was excised, trypsinized and analyzed by mass spectroscopy. Mass spectroscopy was performed on a fee for service basis at the Interdisciplinary Center for Biotechnology Research at the University of Florida. Data was interpreted with Scaffold 4 software from Proteome Software (Portland, Oregon).

### Epitope Mapping

The epitopes of the pNF-H antibodies employed here have already been mapped to the phosphorylated lysine-serine-proline (KSP) sequences in the C-terminal tail region [35]. To map the UCHL1 monoclonals we developed a dot blot assay using a set of 12 amino acid peptides staggered by 5 amino acids based on the human full length sequence, amino acids 1-12, 6-17, 11-22 and so on, a total of 43 peptides. The 12 amino acid peptides in 0.5 mg amounts were dissolved in 200 μL of 50% DMSO/50% distilled water. The peptides were arrayed in a grid pattern on nitrocellulose membranes in 0.5 μL amounts, and the membranes were air dried and blocked for one hour with 5% non fat milk in TBSt. The membranes were then incubated with 1 μg/mL of the relevant antibody and processed with appropriate washing and incubation with goat anti-mouse phosphatase to allow signal visualization with 5-bromo-4-chloro-3-indolyl phosphate/p-nitroblue tetrazolium chloride (BCIP/NBT). To confirm the results from the dot blot assay we developed a competitive ELISA using a set of 20 amino acid peptides based on the human sequence, each peptide overlapping the next by 5 amino acids at each end, so in this case 1-20, 16-35, 31-50 and so on. The 20 amino acid peptides were also in 0.5 mg amounts and each was dissolved in 500 μL of 50% acetonitrile/50% distilled water, so 1 mg/mL. ELISA plates were coated overnight with 0.5 μg/mL of recombinant human UCHL1, blocked in 5% non fat milk in TBSt, and washed on a plate washer with TBSt. In parallel, 100 μL of 1 μg/mL of the relevant antibody was put into each well of another 96 well plate and sequentially 10 μL of each peptide solution added and incubated for 30 minutes at room temperature. All peptide/antibody mixtures were then transferred to the UCHL1 protein coated plate, which was incubated for 1 hour at room temperature. Following three washes in TBSt, incubation with goat anti-mouse alkaline phosphatase secondary, three further washes in TBSt, signal was developed with p-nitrophenyl phosphate (pNPP).

### Statistical analysis

Analysis of data was performed using R and SAS version 9.1 and Figures 2, 3, 4 and 5 were generated using Graphpad Prism 6.0.

## Results

### pNF-H and UCHL1 in control serum samples

We analyzed 20 control blood samples from apparently healthy individuals which were obtained commercially and the levels of UCHL1 detected by the MSD assay were 199±296 ng/mL. One of the individuals had a blood UCHL1 level of 1.33 ng/mL, but even disregarding this individual there was considerable variability in the remainder, five showing no detectable signal whilst the two next highest had signals of 380 and 388 ng/mL. There was no obvious correlation of these blood levels with either age or sex of the individual. We thought these control levels were rather high so we wondered if this was due to background in our assay or the presence of a variable amount of UCHL1 in control blood. Accordingly we selected four control blood samples which we had determined had about average UCHL1 levels, immunodepleted them of IgG and human serum albumin using SpinTrap columns and probed them with our panel of UCHL1 antibodies. Both monoclonal and polyclonal antibodies gave clean and convincing signals at 24 kDa, the size expected for intact UCHL1 (Figure 1A). As shown below, the monoclonal antibodies shown map to distinct epitopes on the UCHL1 molecule. Recombinant human UCHL1 was used as a positive control, giving a strong signal at about 30 kDa, higher than native UCHL1, as expected due to the additional presence of a His tag and other vector derived sequences, 52 amino acids amounting to about 6 kDa. Encouraged by these findings we cut out the 24 kDa region of a Coomassie stained gel from one of these blood samples and performed mass spectroscopy. As shown in Figure 1B, we detected 8 unique peptides derived from human UCHL1, highlighted in yellow, and UCHL1 was identified with 100% confidence.

**Figure 1.**
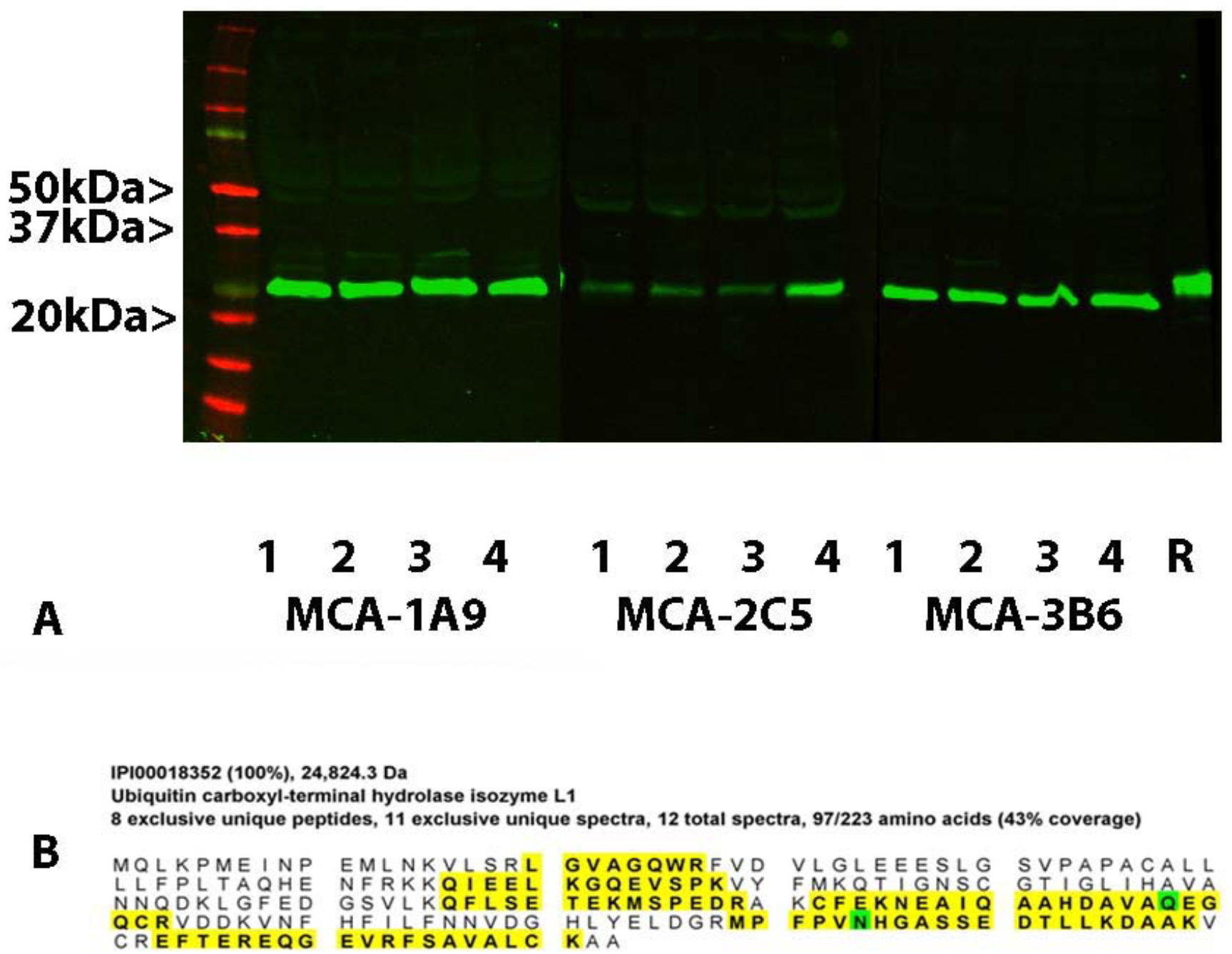
Detection of UCHL1 in control human blood. Panel A shows, in the first lane, molecular weight standards of the indicated molecular size. Lanes labeled 1-4 are western blots of four different control blood samples processed to remove major serum proteins which gave positive MSD assay signals for UCHL1 and which were blotted with UCHL1 monoclonal antibodies. All three antibodies shown give strong signals at the expected 24 kDa molecular weight. The lane labeled R on the far right shows a lane blotted with MCA-3B6 of recombinant human UCHL1, which is about 6 kDa larger in molecular size due to the presence of a His-tag and some other vector derived sequence. Panel B shows mass spectroscopic data from the 24 kDa band from one of the UCHL1 positive blood samples. 8 unique peptides derived from human UCHL1 are identified with high confidence.

**Figure 2.**
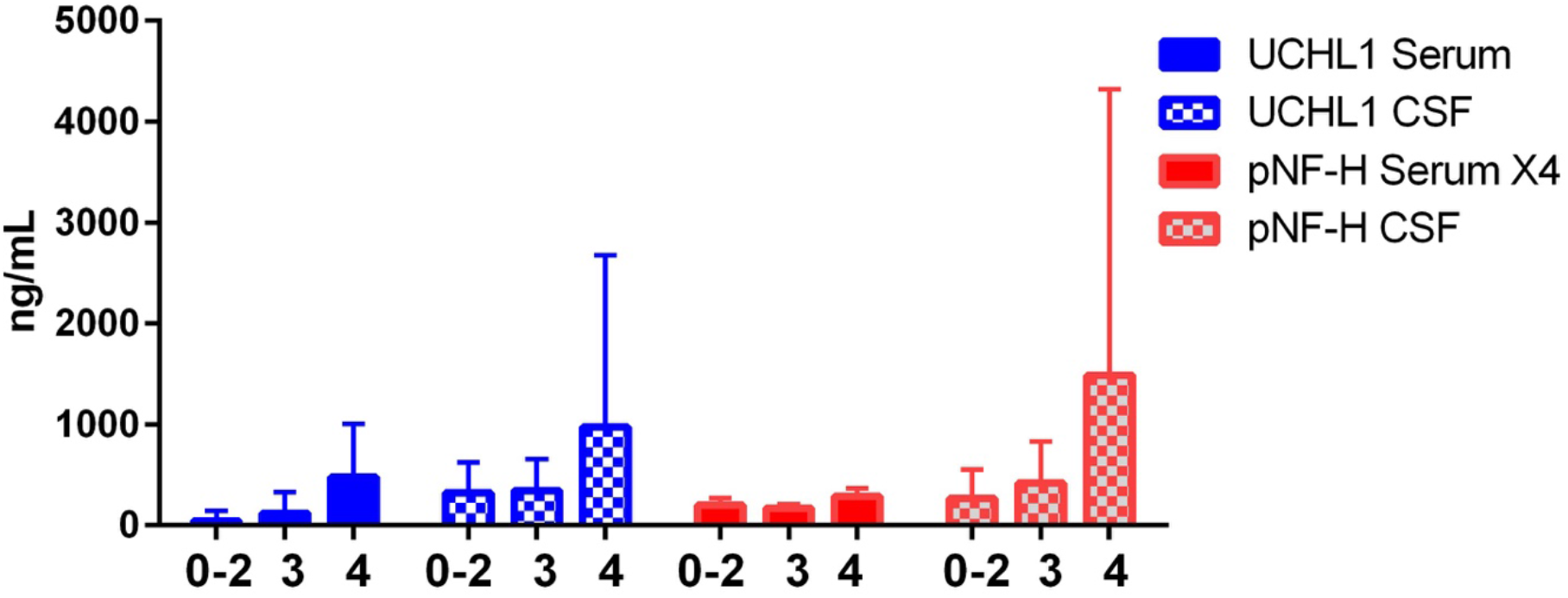
Averages of UCHL1 and pNF-H values of all blood and CSF samples from each patient as a function of patient disposition at discharge. Blood levels of pNF-H were multiplied by four to make them more visible. 0-2 represent the best patient outcome, 3 and 4 are progressively worse.

**Figure 3.**
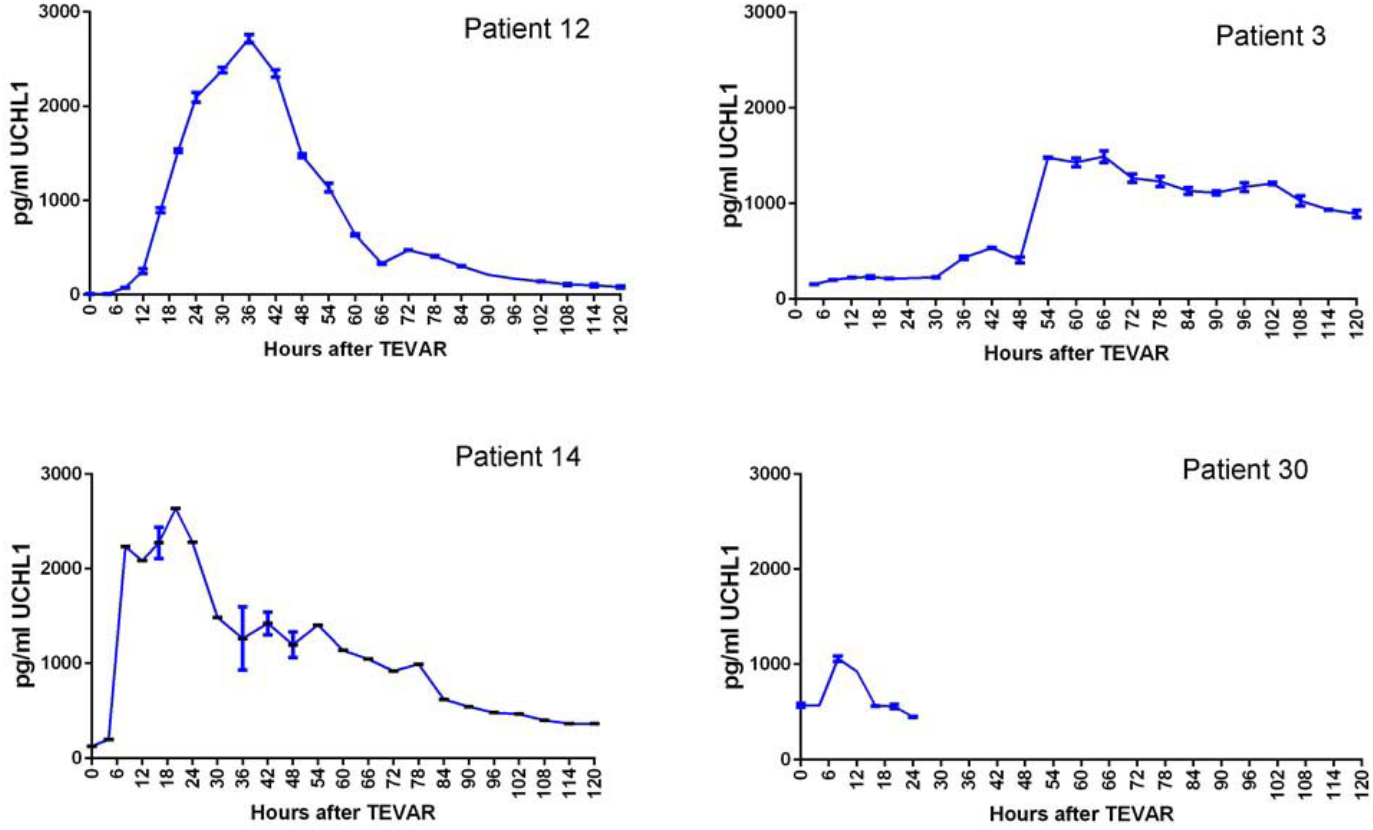
Unusually high blood levels of UCHL1 in 4 TEVAR patients. Error bars are standard deviations. These are the only patients who revealed blood UCHL1 levels over 1 ng/mL at any time in this study, and patients 3, 12 and 14 expired, while patient 30 recovered.

**Figure 4.**
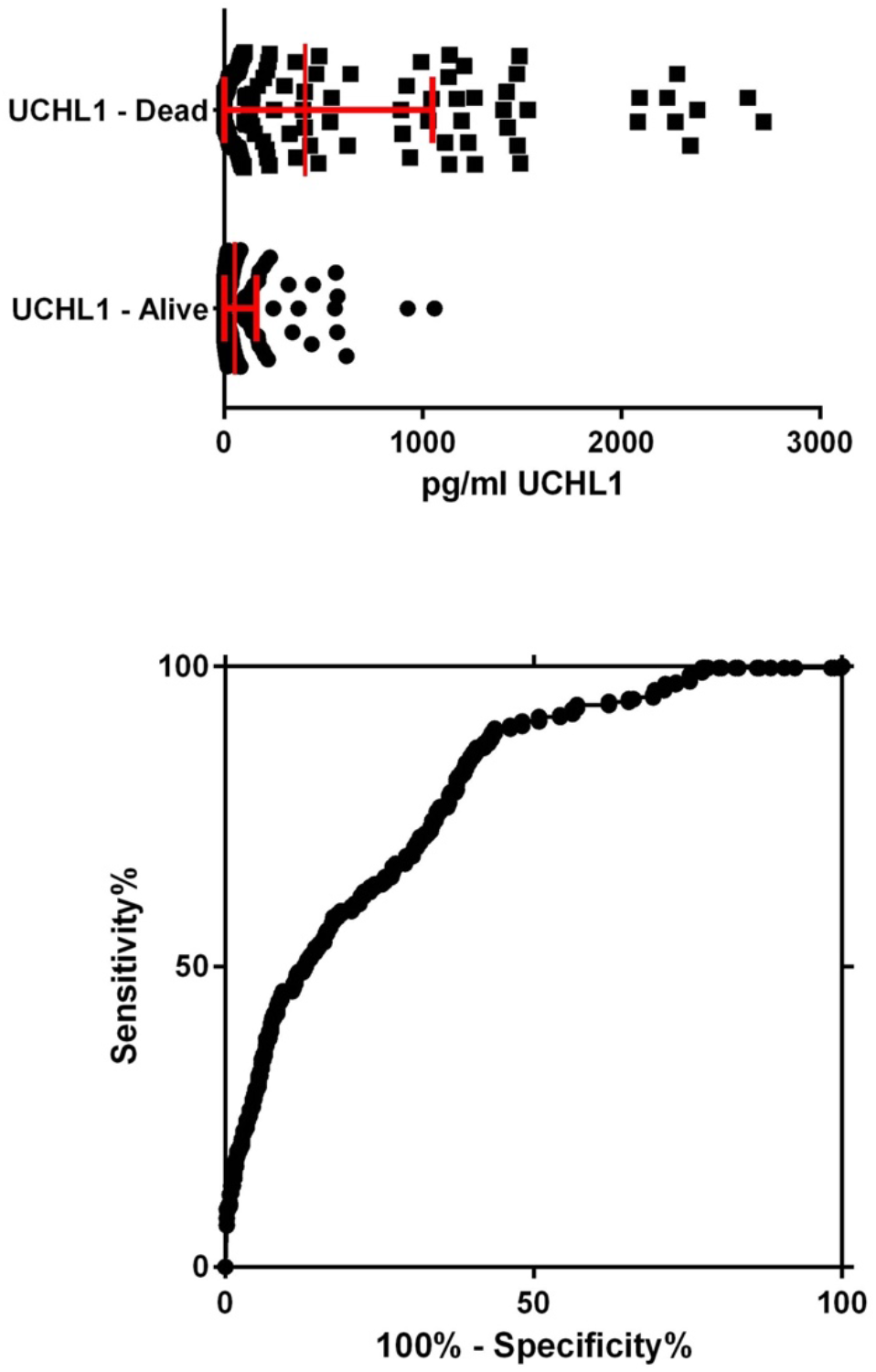
Levels of blood UCHL1 as a function of patient outcome, living or expired, at one year post TEVAR. Top, raw data presented as a dot plot. Clearly high blood levels of UCHL1 predict the worst outcome. Bottom, ROC curve of same data, giving an area under the curve (AUC) of 0.8227.

**Figure 5.**
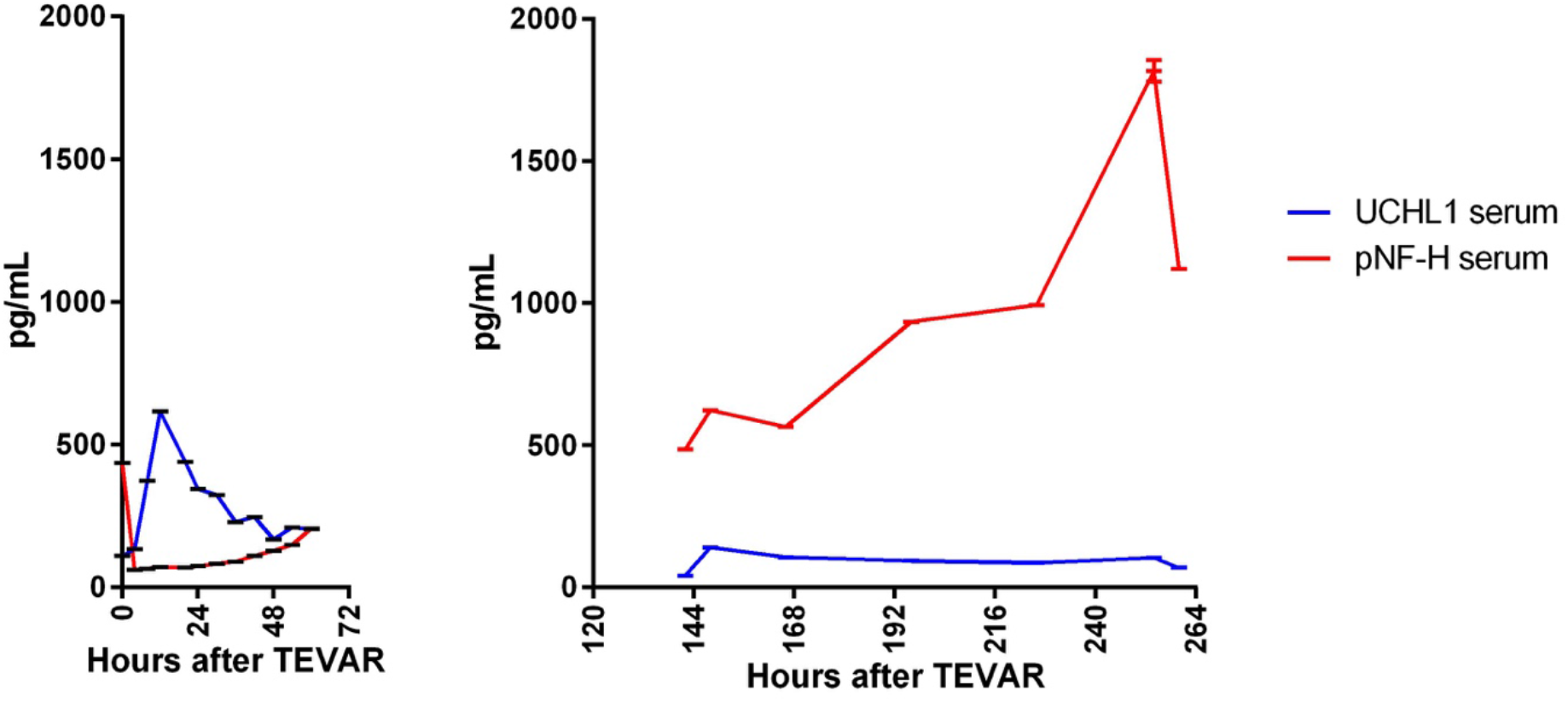
Blood pNF-H and UCHL1 in a single patient who had samples collected over two periods in intensive care out to 11 days after the original operation. The blood pNF-H level slowly increases over the entire period while UCHL1 levels declined.

From several previous studies the baseline levels of pNF-H of healthy control individuals using ELISA has been established to be less than 0.2 ng/mL in plasma and serum and less than 0.8 ng/mL in CSF [16, 18, 37, 38]. Here we establish similar findings using our novel electrochemiluminescent assay, which revealed an average serum pNF-H level of 26.29±29.72 pg/mL using the 20 healthy serum controls. In a previous study using a monoclonal antibody based ELISA we noted that pNF-H levels in control blood samples were low, with a median of 0.17 ng/mL in a cohort of 19 controls [35]. These levels were significantly higher in men compared to women and a weak positive correlation between age and blood pNF-H level was noted (p=0.063). The significance of these findings are not clear given the low sample size, but suggested that some pNF-H may leak into blood normally, possibly in increasing amounts as a function of aging. In the present study we used a different platform, a different set of controls, and different pNF-H detection and capture immunoreagents and found comparable average serum pNF-H levels of 26.3±29.7 ng/mL. Interestingly there was again a weak positive but not statistically significant relationship with age, giving a Pearson correlation coefficient r value of 0.2327, again consistent with a low level of age related pNF-H release into blood. However western blotting of the control human serum samples, unlike UCHL1 as shown here, did not give a convincing signal for full length pNF-H possibly due to the lower apparent protein level, proteolysis or other processing and/or various technical issues. We therefore currently have no evidence independent of ELISA type assays showing that pNF-H is present in normal human blood.

### General overview of TEVAR patient results

We analyzed a total of 536 serum, 295 CSF and 108 urine samples from 50 TEVAR patients. In a preliminary analysis we simply averaged the levels of the two biomarkers in blood and CSF as a function of patient disposition at discharge. Both biomarkers in both biofluids showed elevated levels in patients with poorer dispositions (Figure 2). UCHL1 levels were higher than pNF-H levels and patients with the best outcomes had close to control levels of both proteins. Comparison of survivors to those who expired in terms of serum UCHL1 gave a p value of less than 0.0001. For the same comparison pNF-H serum levels were not statistically significant, but CSF levels of both proteins indicated p values less than 0.0001. Levels of the two proteins in the more limited set of urine samples also showed increased levels in patients with the worse outcomes, though, possibly because of the more limited number of patients, there was no obvious correlation with outcome.

### Trajectory of change in biomarker levels over time

We used mixed effects linear regression models to characterize the trajectory of blood and CSF biomarker levels across time and to estimate the effects of SCI and the right leg exam (RLE) on these levels in 50 patients. The RLE scale is used to estimate the degree of patient SCI. Log values of serum UCHL1 declined by 0.008 per hour on average (95% CI= [−0.015, −0.001]. p=0.0346), while for serum pNF-H log values rose by 0.008 per hour on average (95% CI= [−0.001, − 0.012]. p=0.003). For CSF levels, the average level of UCHL1 did not vary significantly over time over the whole group, although patients with SCI had 0.66 higher levels on average (95% CI= [−0.101, −1.43]. p=0.087). The log pNF-H CSF levels were significantly higher in SCI patients than the rest of the cohort (p=0.0483).

A mixed-effects linear regression analysis of urine samples from 16 patients showed that log(UCHL1) levels increased 0.033 per hour post surgery and that SCI patients showed an average of 1.2 units higher level than non-SCI patients. There was also a significant relationship between log(UCHL1) levels and the RLE values, with an increase in log(UCHL1) of 0.81 for each increase in RLE score (CI=[0.017, 1.60], p=.045). Urine levels of pNF-H showed no statistically significant change over time.

### Details of Biomarker Release Profiles

Inspection of the profiles of biomarker release of TEVAR patients revealed many interesting observations. A notable feature seen in the case of four patients was marked sustained release of UCHL1 peaking at over 1 ng/mL into serum in the hours after the TEVAR operation (top, Figure 3). Interestingly, patients 3 and 12 were diagnosed with renal failure, while patient 14 was diagnosed as having acute renal failure. These were the only patients with renal failure in the cohort, suggesting a strong relationship between elevated blood UCHL1 levels and renal compromise. These three patients also had the worst outcomes, expiring at 37 days, 100 days and 45 days after TEVAR respectively. It is of note that the same three patients showed clinical symptoms of spinal cord ischemia (SCI), although they were not unique in this respect, since 5 other patients showed SCI symptoms without showing such marked UCHL1 release into blood. Patient 30 was the only other patient revealing more than 1 ng/mL blood UCHL1 at any time following the operation in this group, which we saw in a single blood sample (Figure 3). This patient had the lowest peak over 1 ng/mL and average levels of blood UCHL1, and it is notable that the patient was still living a year after TEVAR.

We performed receiver operating characteristic (ROC) analysis of the all blood UCHL1 levels measured in this study, dichotomized for outcome, either dead or alive within 12 months of a TEVAR procedure, and the data is shown in Figure 4. The area under the curve (AUC) is 0.8227, suggestive of some diagnostic potential. We can conclude that high levels of blood UCHL1 detected at any time in the first 5 days following a TEVAR procedure are associated with the poorest patient outcome.

There was a general correlation between blood pNF-H levels and outcome. One individual, patient 34, underwent treatment for an unusually long period, and as a result samples of blood and CSF were taken over two time periods, the second being out to 11 days following the original operation (Figure 5, left panel 0-60 hours, right panel 144-264 hours). As shown this patient also showed a modest peak of release of UCHL1 in the first few hours post-TEVAR, though the amplitude of this peak was lower than seen in patients 3, 12, 14 and 30. Interestingly, the levels of pNF-H persistently increased not only over the first 5 day period but also continued to increase well beyond this time. This patient, like 3, 12, 14 and 30, showed symptoms of SCI and also did not recover fully.

### Blood samples taken at 30 days after TEVAR

Blood samples were taken from 9 patients at the 30 day follow up. An interesting feature of this was strong signals for pNF-H, ranging from 640 pg/mL to 998 pg/ml in 3 of these patients. Two of the patients with strong pNF-H signals were among those with less good outcomes, ambulating only with help at discharge and follow up, while the third was ambulating normally at discharge. 5 of the patients with a normal blood pNF-H signal were ambulating only with help at discharge, while the remaining 1 patient was not. These findings are extremely preliminary given the small sample size but suggest that 30 day release of pNF-H is correlated with poorer outcome. All of these 30 day blood samples gave very low UCHL1 signals within the range established here, from undetectable to 102 ng/mL.

### CSF samples

In line with previous studies the levels of CSF, both UCHL1 and pNF-H levels were much higher than seen in blood, the highest levels being over 9 ng/mL for UCHL1 and almost 19 ng/mL for pNF-H. As with the blood values, higher levels of both proteins were associated with poorer outcomes. CSF levels above 5 ng/mL sampled at any time following TEVAR for either biomarker predicted death within the study period with 100% sensitivity. However lower CSF levels were less predictive than blood UCHL1 levels giving AUCs of 0.67 for UCHL1 and 0.66 for pNF-H.

### Urine samples

We were surprised to find strong MSD assay signals for UCHL1 in some urine samples. In most cases these large signals were observed in samples taken 12-48 hours following TEVAR, suggesting that the release of this protein into urine was secondary to the TEVAR operation (Figure 6). We saw much weaker pNF-H MSD assay signals in these samples, the pNF-H levels ranging from undetectable to a maximum of 32 pg/mL. Interestingly, two out of four patients who had the highest urine levels of UCHL1 had suffered from strokes. Patient 34, the single one from whom we obtained blood, CSF and urine samples out to 154 hours post TEVAR, also revealed high levels of UCHL1 in urine over the time period from 60-154 hours. This patient revealed one urine sample with over 2 ng/mL UCHL1 and also showed consistent pNF-H signals over 10 pg/mL, peaking at 30.1 pg/mL.

**Figure 6.**
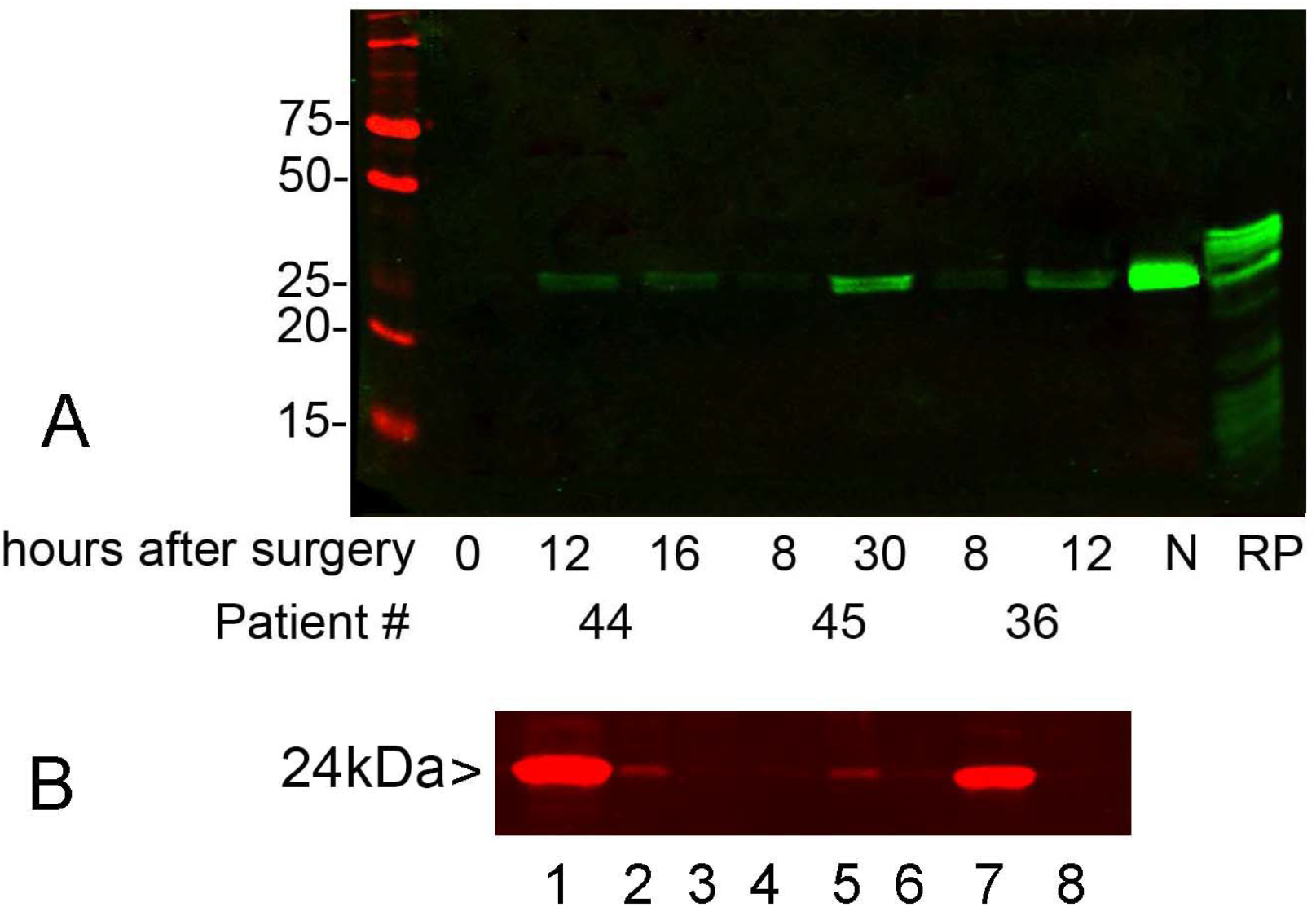
Panel A shows detection of UCHL1 in urine by western blotting. Urine samples from the three indicated patients were taken at the times also indicated and probed with MCA-BH7 monoclonal antibody to UCHL1. N is a sample of urine taken from a neonatal hypoxic ischemic encephalopathy patient from a previous study which showed a very strong UCHL1 signal by ELISA [39]. RP is a preparation of recombinant UCHL1, which is an about 6 kDa larger in apparent molecular size due to the presence of an N-terminal His tag and other vector derived sequence. This preparation had a series of lower molecular weight bands due to partial proteolysis during preparation. Samples in leftmost lane are molecular weight standards of size indicated in kilodaltons. Panel B shows the relevant region of a western blot of equivalent protein levels of crude extracts of various mouse tissues probed with rabbit polyclonal antibody to UCHL1. 1 = brain, 2 = heart, 3 = liver, 4 = spleen, 5 = kidney and 6 = lung. Lanes 7 and 8 were crude extracts of HEK293 and HeLa cells respectively. Significant levels of UCHL1 can be readily detected in both heart and kidney extracts and on longer blot exposure all tissues tested showed a UCHL1 positive band. A similar probe of rat tissues gave identical results. HEK293 cells are known to express UCHL1 and other neuronal proteins [58], while HeLa cells are negative for UCHL1 and are non-neuronal in origin.

To determine whether urine levels and serum levels of UCHL1 were associated, we ran a linear mixed model with log(serum level) as outcome and log(urine level) as predictor for the 16 patients for which we had the appropriate data. The result was p=0.035, indicating that urine and serum levels are significantly related. The mean correlation, weighted by the number of observations was 0.27. We noted that the association between serum and urine is highly patient-specific. For some the correlation is nearly perfect, while for others, there was little correlation. We did a similar analysis for blood pNF-H versus urine pNF-H and found no significant relationship.

We also ran a linear mixed model with log(urine UCHL1 level) as outcome and log(urine pNF-H level) as predictor. The result was p=0.0007, indicating that these levels are very significantly related, with a correlation of O.19. So, although the pNF-H levels were orders of magnitude lower than the UCHL1 levels, the two proteins appear to be leaking into urine in a parallel fashion.

### Western blotting of urine and rat tissues

To confirm the presence of UCHL1 in urine we performed western blotting of a subset of the UCHL1 positive and negative urine samples (Figure 6A). Urine from patient 44 was essentially negative for UCHL1 at time zero, and levels became readily detectable at 12 and 16 hours. As shown, a protein band at about 24 kDa is scarcely visible in the time zero lane but becomes obvious in the 12 and 16 hour lanes. Consistent with these findings the MSD assay detected 43 pg/ml, 953 pg/ml and 818 pg/mL of UCHL1 in samples corresponding to those in those three lanes respectively. The MSD data were similarly in line with the two samples from patients 45 (29 pg/mL, 1.49 ng/mL respectively), and 36 (160 pg/mL and 821 pg/mL respectively). We included a urine sample taken as part of a previous study of neonates suffering from hypoxic ischemic encephalopathy [39]. This sample had shown a very strong MSD signal, over 5 ng/mL which saturated our assay, and which, as shown, had the strongest western blot band intensity also. The blot illustrated was performed using MCA-BH7, a monoclonal antibody to UCHL1 used as the capture reagent in the MSD assay used here. Identical results were obtained with other UCHL1 monoclonal antibodies and our affinity purified rabbit antibody to UCHL1 (not shown).

As discussed in more detail below there is immunohistological evidence for the presence of UCHL1 in tissues other than brain, with kidney being of particular interest given the presence of this protein in urine. Accordingly we probed several rat and mouse tissue extracts with UCHL1 antibody and noted, as expected, a prominent signal in brain extract, but also clear signals in both cardiac and kidney extracts (lanes 1, 5 and 7 respectively, Figure 6B). Very faint bands were visible on longer blot exposure in other tissues.

### Antibody Epitope Mapping

The UCHL1 monoclonal antibody MCA-BH7, the capture reagent in the ELISA and MSD assays described here, is available commercially and has been widely used. This antibody gave a strong signal in our dot blot assay showing strong binding with the peptide WRFVDVLGLEEE, human UCHL1 amino acids 26-37. A disadvantage to this assay was that one other peptide gave a very strong signal with all four UCHL1 monoclonal antibodies tested. We hypothesized that this peptide must bind strongly to some region of the mouse IgG unrelated to the antigen binding sites. We are studying this further as it may represent the basis for the development of a novel IgG recognition reagent. To obtain more hopefully confirmatory data we developed a competitive ELISA assay using 20 amino acid peptides each overlapping the next by 5 amino acids. The peptide VLSRLGVAGQWRFVDVLGLE, amino acids 16-35 of the human sequence, strongly and reproducibly inhibited binding of MCA-BH7 to recombinant UCHL1. Since the preceding and following 20 amino acid peptides did not interfere with binding we conclude that the central sequence, GVAGQWRFVD, amino acids 21-30, is key for antibody binding. The finding with the 12 amino acid sequence, WRFVDVLGLEEE, is consistent with this, focusing attention on the WRFVD sequence present in both peptides. This is amino acids 26-30 which corresponds to a short β-strand close to the UCHL1 N-terminus, see https://www.rcsb.org/3d-view/2ETL/1.

The antibody MCA-2C5 binds the peptide KGQEVSPKVYFM in the dot blot assay, amino acids 71-82. The binding of the MCA-2C5 antibody was inhibited strongly by the peptide NFRKKQIEELKGQEVSPKVY, amino acids 61-80 in the competitive ELISA assay. Again the previous and following 20 amino acid peptides had no obvious effect, focusing attention on the peptide QIEELKGQEV, amino acids 66-75. We conclude that the peptide KGQEV, present in both peptides, is the core of the MCA-2A5 epitope. This is amino acids 71-75 of the human sequence and corresponds to a turn at end of a long α-helix, see https://www.rcsb.org/3d-view/2ETL/1.

As is often the case with these types of experiment, no clear dot blot or competitive ELISA result was seen with the two other UCHL1 monoclonals tested, MCA-1A4 and MCA-3B6. This is likely due to these antibodies being dependent on non-contiguous or highly conformation dependent epitopes. However we can conclude that these four UCHL1 antibodies must bind a minimum of three different epitopes, building overwhelming confidence in our western blotting results.

## Discussion

As far as we are aware the present study is the first to look at blood, CSF and urine samples from a cohort of TEVAR patients using assays for either pNF-H or UCHL1. We found that both proteins could be detected at elevated levels in all three biofluids in many patients and that higher levels were in general predictive of a worse outcome (Figure 2). Particularly striking were peaks of UCHL1 release into the blood seen in four of the patients after TEVAR (Figure 3). These peaks appeared to initiate suddenly and were persistent, slowly reducing in level over several days. The sudden initiation suggested that the peaks of release were the result of acute events. Since UCHL1 is concentrated in neuronal perikarya, the origin of this protein in blood would normally be thought to be due to gray matter compromise, though we will discuss this issue in more detail below. It is striking that the majority of the individuals with blood levels over 1 ng/mL at any time post operation were destined to expire. This suggests that detection of such high levels of UCHL1 in the blood after a TEVAR is related to serious events secondary to the operation predictive of a poor outcome, and it is striking that this prediction was fulfilled several weeks or months after the operation. This conclusion is in line with our previous studies which showed sudden release of large amounts of UCHL1 into blood following traumatic brain injury [40] and also following hypoxic ischemic encephalopathy (HIE) both in neonates [41] and in foals [42]. The number of patients who showed these phenomena, 4 out of 50, is about the number expected for patients suffering from SCI in a typical TEVAR population. Patients 3, 12 and 14 showed symptoms of leg weakness, which is used as a diagnostic criteria for SCI. Taken together these findings suggest that profound CNS compromise may release readily measurable amount of UCHL1 into blood, and supports our general conclusion that elevated blood UCHL1 is a useful marker of acute CNS compromise. A tentative conclusion from the present study is that blood levels of UCHL1 above 1.0 ng/mL detected at any time after TEVAR are associated with SCI and with the worst outcome in these patients. Further study will be needed to translate this finding into practice.

With any antibody based assay for a blood biomarker there is always the issue of whether a signal seen in control sera is due to a small amount of the analyte being present or is due to background in the assay. The results with control blood samples probed for UCHL1 with our MSD assay showed signals averaging about 0.2 ng/mL but with great variability. We found unequivocal evidence for the presence of intact UCHL1 in control serum samples by western blotting and mass spectroscopy, confirming data obtained with the MSD assay. The great variability in the control samples suggests that there may be some diurnal, stress related or other periodic pattern of release, and further examination of this issue is of course of some interest. The origins and mechanisms of UCHL1 release into blood are of interest. Exosomes are present in blood and some of these may be derived from nerve endings expected to be rich in UCHL1 [43]. Exosomes containing UCHL1 could also originate from neuroendocrine cells or some of the other non-CNS cell types known to express this protein. Many previous reviews have made the claim that UCHL1 is expressed only in brain [44]. However UCHL1 is a major protein of spinal cord and peripheral ganglia. It is also a defining component of nerve fibers and is found in neuroendocrine cells, both of which may be outside the CNS. Antibodies to Pgp9.5, now identified as UCHL1, are used to identify nerve fibers in skin and other tissues [32]. Since all tissues contain nerve fibers, UCHL1 will be ubiquitously expressed in at least low levels. As shown here, all rat and mouse tissues probed with UCHL1 antibody showed a weak positive band and both kidney and cardiac tissue showed a more substantial band. In agreement with our findings, inspection of data in the Human Protein Atlas (http://www.proteinatlas.org/ENSG00000154277-UCHL1/tissue) shows high protein and mRNA expression for UCHL1 in the central nervous system as expected. The atlas also shows high protein levels in the pancreas and kidney, while lower levels were seen in soft tissue, testes, ovaries and colon. In the kidney older immunocytochemical studies with UCHL1 antibodies show positive staining in the human nephron epithelia [45] and in the rat [44, 46]. There are also several different damage and disease states associated with release of non-CNS origin UCHL1 into blood. For example UCHL1 is an abundant cytoplasmic protein of pancreatic β-cells, and injection of streptozotocin into rats destroys these cells and results in the rapid release of UCHL1 into the blood over a time period of 2-6 hours [47]. In some animals peak UCHL1 levels were reported to be over 20 pmol/L, which translates to 0.48 ng/mL. This raises the issue of whether blood UCHL1 release might be seen in diabetic or pre-diabetic patients. A large separate literature has focused on UCHL1 as an abundant protein of certain colorectal, pancreatic, parathyroid and lung cancers with higher expression often being correlated with poorer outcome [48–50]. It is well known that the contents of cancer cells are released into blood and other bodily fluids. All this work raises serious questions about the origin of blood UCHL1 in healthy individuals and also polytrauma patients, especially if the kidneys or heart are damaged, the patient may be prediabetic or have a tumor. Clearly much work will have to be performed before UCHL1 blood levels can be reliably used as a general CNS damage or disease biomarker.

In this particular patient group levels of pNF-H proved to be less generally predictive than UCHL1 levels. As with several of our previous studies we saw slow release of pNF-H into blood in some patients over 5 days post TEVAR, and the patients showing this generally had worse outcomes. We analyzed a more extensive time period for patient 34 who revealed a persistent release pattern (Figure 5). The marked elevations in the level of pNF-H in some of the blood samples taken 30 days after TEVAR are in line with this suggestion also. We made somewhat comparable findings in a previous study of aneurysmal subarachnoid hemorrhage (ASAH) patients, although these blood samples were taken over 14 days post stroke and surgery so that the slow persistent increase in blood levels of this protein could be better visualized [16]. In these ASAH cases the level of blood pNF-H was strongly correlated with ASAH patient outcome, higher levels predicting poorer outcomes.

Several other studies suggest that blood pNF-H levels provide a useful biomarker of ongoing axonal loss in multiple sclerosis [51], amyotrophic lateral sclerosis [35, 38] and other neurodegenerative states. We have proposed that this release reflects ongoing axonal loss secondary to a variety of different CNS compromises and several other groups have made similar findings [28]. The small number of samples collected after 5 days in the present study do not allow us to firmly make the conclusion that such higher levels are related to poorer outcome in TEVAR patients, but inspection of available data suggests that this is the case. In summary, the present findings are in line with our proposal that persistent increases in the blood pNF-H level days and even weeks after a CNS compromise is a novel means of assessing ongoing chronic and secondary axonal loss.

As expected CSF levels both of pNF-H and UCHL1 were much higher than blood levels, and higher levels of both were associated with poorer outcomes. A somewhat surprising finding was the presence of high levels of UCHL1 in some of the urine samples taken. The levels ranged from 0 to over 2 ng/ml for UCHL1 and 0 to 31 pg/ml for pNF-H. Although the levels of pNF-H were much lower they were significantly associated with the UCHL1 levels, suggesting that both were entering urine by a single mechanism. We were also able to confirm the presence of UCHL1 in urine by western blotting, showing for the first time that urine may contain intact full length UCHL1 (Figure 6A). We also show the presence of readily detectable levels of UCHL1 in kidney and cardiac tissues (Figure 6B).

Neither UCHL1 nor pNF-H were found in an initial screen of the human urinary proteome which however found evidence for the presence of 1,500 other proteins [52]. The present study revealed substantial amounts of UCHL1 and pNF-H only in the urine of severely compromised individuals, and we could not detect UCHL1 or pNF-H in most urine samples studied here or in samples obtained from several normal healthy individuals (not shown). We also noted even larger amounts of UCHL1 in urine from certain neonatal HIE patients, but not in control neonatal urine samples. UCHL1 rich urine samples in these neonates was associated with very poor outcomes and also with renal failure. The origin of these proteins in urine is unknown. Although the level of UCHL1 in blood and urine appeared to related, the level in urine was generally much higher, suggesting it was not simply leaking from blood. Possibly some UCHL1 is leaking directly from the kidney, and it is notable that three out of the four patients shown in Figure 3 had renal failure. The much lower pNF-H urine signal was however significantly associated with the UCHL1 urine signal. This pNF-H may also be of kidney origin; The kidney is innervated by axons which express pNF-H and UCHL1. Accordingly a serious compromise to the kidney would be expected to result in the release of small amounts of pNF-H and UCHL1 from endogenous nerve fibers. However the majority of the UCHL1 detected may derive directly from the kidney epithelium in individuals with renal failure. These findings may provide a fruitful avenue for future research and may prove to be of clinical utility. Urine samples are of course one of the least invasive sources for biomarker measurement.

A few previous studies have examined levels of either pNF-H or UCHL1 following traumatic SCI. Yokobori et al. authored a review on potential biomarkers of SCI which includes some data suggesting that UCHL1 may be released into CSF of rodents and patients in elevated amounts following SCI [53]. However in neither case were the levels of blood UCHL1 addressed and the patient data is extremely preliminary, with samples from only a single patient being shown. Hayakawa et al. showed that pNF-H levels were significantly higher in patients with American Spinal Injury Association Impairment Scale (AIS) scale A, the most serious, compared to those with AIS scale C, AIS D and AIS E over the time period from 6 hours to 21 days post injury [54]. These authors noted a slow persistent increase in blood levels of pNF-H as seen here. Ahadi et al. looked at blood levels of pNF-H in 35 traumatic SCI patients, 16 of which were AIS A grade, 24-48 hours after injury [55]. They found very significant increases in the average level of blood pNF-H compared to uninjured controls within 24 hours of injury. Kuhle et al. studied blood levels of a different neurofilament subunit, NF-L in the blood of a cohort of patients with spinal cord injury [56]. They found that higher levels of blood NF-L were found in patients with complete spinal cord section compared to those with incomplete section and both were significantly higher than controls. Samples were taken every 12 hours out to 7 days and levels of NF-L persistently increased over that time a finding similar to that we made on ASAH patients [16] and in the present study. Interestingly the blood level of NF-L were lower and approached significance in patients treated with minocycline, an anti-inflammatory drug, which suggests that blood neurofilament levels might ultimately be used to monitor recovery and response to therapy. More recently Merisson et al. looked at blood levels of NF-L and total MAP-tau in CSF of a small cohort of 10 patients following TEVAR, looking for biomarker evidence of SCI [57]. Both MAP-tau and NF-L are highly concentrated in axons, so similar results to our data on pNF-H might be expected. In fact they found that both proteins were increased in level in the CSF of patients with secondary spinal cord injury though a few hours later than symptoms were detected. Data on plasma or serum samples were not presented in this study.

We here describe novel assays developed for the electrochemiluminescence based MSD platform which have several advantages over regular ELISA. These assays have a greater dynamic range, slightly greater sensitivity and requires a significantly smaller sample volume. With further development it should be possible to develop an MSD based assay capable of monitoring UCHL1, pNF-H and other analytes in a single well, with considerable savings in sample utilization, time and effort. Finally we further epitope mapped two of our current panel of monoclonal antibodies using a peptide dot blot assay in combination with a competitive peptide ELISA, an approach which may be useful for other researchers. The widely used mouse monoclonal antibody to UCHL1 MCA-BH7 is therefore unusually well characterized, since we also recently characterized the K_D_ and other kinetic properties of this antibody (see https://encorbio.com/product/MCA-BH7).

In summary, this report describes several novel and potentially clinically relevant findings with respect to TEVAR patients and extends further the utility of UCHL1 and pNF-H assays. The presence of UCHL1 in control blood samples, in kidney tissues and in urine samples all raise questions about the feasibility of using this biomarker for clinical purposes. However, in future studies it may be possible to monitor patient response to the TEVAR operation, progression, occurrence of secondary problems, response to therapy and predict patient outcome using simple blood, CSF and urine based assays. In particular, the measurement of UCHL1 in urine appears to be a fruitful avenue for further study.

## Acknowledgments

Supported by grants from Brain and Spinal Cord Injury Trust Fund of the state of Florida, the NIH and private funding from EnCor Biotechnology Inc.

